# A female-specific role for Calcitonin Gene-Related Peptide (CGRP) in rodent pain models

**DOI:** 10.1101/2021.06.02.446716

**Authors:** Candler Paige, Isabel Plasencia-Fernandez, Moeno Kume, Melina Papalampropoulou-Tsiridou, Louis-Etienne Lorenzo, Galo L. Mejia, Christopher Driskill, Francesco Ferrini, Andrew L. Feldhaus, Leon F. Garcia-Martinez, Armen N. Akopian, Yves De Koninck, Gregory Dussor, Theodore J. Price

## Abstract

We aimed to investigate a potentially sexually dimorphic role of Calcitonin Gene-Related Peptide (CGRP) in mouse and rat models of pain. Based on findings in migraine where CGRP has a preferential pain-promoting effect in female rodents, we hypothesized that CGRP antagonists and antibodies would attenuate pain sensitization more efficaciously in female than male mice and rats. In hyperalgesic priming induced by activation of interleukin 6 (IL-6) signaling, CGRP receptor antagonists, olcegepant and CGRP_8-37_, both given intrathecally, blocked and reversed hyperalgesic priming only in females. A monoclonal antibody against CGRP, given systemically, blocked priming specifically in female rodents but failed to reverse it. In the spared nerve injury (SNI) model, there was a transient effect of both CGRP antagonists, given intrathecally, on mechanical hypersensitivity in female mice only. Consistent with these findings, intrathecally applied CGRP caused a long-lasting, dose-dependent mechanical hypersensitivity in female mice but more transient effects in males. This CGRP-induced mechanical hypersensitivity was reversed by the KCC2 activator, CLP257 suggesting a role for anionic plasticity in the dorsal horn in the pain-promoting effects of CGRP in females. In spinal dorsal horn slices, CGRP shifted GABA_A_ reversal potentials to significantly more positive values but, again, only in female mice. Therefore, CGRP may regulate KCC2 expression and/or activity specifically in females. However, KCC2 hypofunction promotes mechanical pain hypersensitivity in both sexes because CLP257 alleviated hyperalgesic priming in male and female mice. We conclude that CGRP promotes pain plasticity in female mice, but has a limited impact in male mice.

**Significance Statement:** The majority of patients impacted by chronic pain are women. Mechanistic studies in rodents are creating a clear picture that molecular events promoting chronic pain are different in male and female animals. Far more is known about chronic pain mechanisms in male animals. We sought to build on recent evidence showing that CGRP is a more potent and efficacious promoter of headache pain in female than in male rodents. To test this, we used hyperalgesic priming and the spared nerve injury (SNI) neuropathic pain models in mice. Our findings show a clear sex dimorphism wherein CGRP promotes pain in female but not male mice. Our work suggests that CGRP antagonists could be tested for efficacy in women for a broader variety of pain conditions.

## Introduction

Pathways that mediate the development and maintenance of chronic pain are increasingly recognized as distinct in males and females (Mogil, 2020). Underlying mechanisms governing these sex differences are now becoming clear. In males, it has been demonstrated that microglia and macrophage activation contribute strongly to the development of chronic pain, because either blocking microglia activation or depleting animals of microglia can prevent the development of mechanical hypersensitivity after injury in mice (Sorge et al., 2015; Taves et al., 2016; Echeverry et al., 2017; Mapplebeck et al., 2018; Paige et al., 2018; Yu et al., 2020). In females, depletion of microglia does not reverse nerve injury-induced hypersensitivity and it appears that T-cells may play a critical role in promoting chronic pain (Sorge et al., 2015), although there is also evidence that T-cells promote pain resolution in both sexes (Krukowski et al., 2016; Laumet et al., 2019). Another female-specific pain promoting mechanism is prolactin signaling, which sensitizes female nociceptors through a direct action on the prolactin receptor (Patil et al., 2013; Patil et al., 2019b; Paige et al., 2020). Two mechanisms could underlie this sex difference in responses to prolactin, a sexually dimorphic expression pattern for the prolactin receptor in subtypes of dorsal root ganglia (DRG) neurons (Patil et al., 2019a), and/or a female-specific translation of the prolactin receptor at central and peripheral terminals of nociceptors (Patil et al., 2019b; Paige et al., 2020).

One of the most striking sex differences in medicine is the far greater incidence of migraine headache in women than in men (Ashina et al., 2021). A number of new therapeutics for migraine have recently been approved. These new drugs target calcitonin gene-related peptide (CGRP) by sequestering the peptide with an antibody or blocking the receptor with a small molecule antagonist or a function blocking antibody (Dodick et al., 2014; Moreno-Ajona et al., 2020). Work aiming to better understand the mechanisms through which CGRP acts in migraine headache demonstrated a dramatic left-ward shift in the dose-dependent effects of CGRP in promoting pain when applied to the dura of female mice (Avona et al., 2019). Other studies have also demonstrated a sex difference in the expression of CGRP receptor components in the trigeminal nucleus (Ji et al., 2019). However, the pain promoting effects of CGRP are not limited to the trigeminal region. Previous experiments demonstrate that CGRP anti-serum given intrathecally increases nociceptive thresholds in a rat model of arthritis (Kuraishi et al., 1988). Moreover, CGRP applied to the dorsal horn induces signaling that increases synaptic efficacy suggesting a direct action on dorsal horn neurons (Sun et al., 2004). To our knowledge, potential sex differences in the effects of CGRP on the dorsal root ganglion (DRG) or spinal dorsal horn have not been assessed.

Anionic plasticity is an important contributor to chronic pain (Kaila et al., 2014; Price and Prescott, 2015; Lorenzo et al., 2020). Changes in Cl^−^ gradients in dorsal horn neurons can lead to decreased inhibitory efficacy enabling non-noxious stimuli to gain access to the ascending nociceptive pathway (Coull et al., 2003; Coull et al., 2005; Keller et al., 2007; Ferrini et al., 2013). This is an important cause of mechanical allodynia, a prominent feature of pain after injury. Decreased expression or function of the Cl^−^ extrusion transporter, KCC2, is the best understood mechanism for anionic plasticity in the dorsal horn (Coull et al., 2003; Miletic and Miletic, 2008; Asiedu et al., 2012; Ferrini et al., 2013; Li et al., 2016; Dedek et al., 2019; Ferrini et al., 2020; Locke et al., 2020; Lorenzo et al., 2020). While previous studies have demonstrated a key role for KCC2 in mechanical allodynia after injury in both male and female rodents (Mapplebeck et al., 2019), the brain-derived neurotrophic factor (BDNF) microglia signaling mechanism promoting neuropathic pain is engaged only in male mice (Sorge et al., 2015). BDNF promotes hyperalgesic priming in the body only in male mice (Moy et al., 2019) but regulates priming in the cephalic region in both sexes (Burgos-Vega et al., 2016). Given this literature, a goal of our work was to determine whether CGRP may influence anionic plasticity in the spinal dorsal horn in a sex-specific fashion.

We show that 2 structurally distinct CGRP antagonists block and reverse the development of mechanical hypersensitivity specifically in females in hyperalgesic priming, incision, and spared-nerve injury (SNI) - induced neuropathic pain. Likewise, CGRP monoclonal antibodies (mAb) block the establishment of hyperalgesic priming in female but not male rats and mice. Intrathecal (I.T.) CGRP causes increased and temporally prolonged hindpaw hypersensitivity in female mice when compared to male mice. This effect is reversed by I.T. CLP257, a KCC2 activator (Gagnon et al., 2013), suggesting a link between CGRP signaling and KCC2 in the dorsal horn. In direct support of this hypothesis, CGRP applied to the spinal cord causes a depolarization of the GABA_A_ receptor reversal potential (E_GABA_) in female but not male mice. This effect appears to be linked to a change in KCC2 function, but not trafficking. Our findings point to a previously unexplored central role of CGRP in promoting pain in female rodents.

## Methods

### Animals

All animal procedures were approved by the Institutional Animal Care and Use Committee at the University of Texas at Dallas and the Canadian Council on Animal Care and the committee for animal protection of Université Laval (CPAUL). Both male and female animals were used in all experiments unless otherwise noted. Experimenters were blinded to each experimental group for all experiments. Animals used per group are noted in each figure. For mouse behavioral experiments, all animals were bred in the Animal Research Facility at the University of Texas at Dallas and were 7 – 12 weeks old at the time of the experiment. Each group was either Swiss Webster or C57BL6. Animals were housed in same-sex groups of 2-4 on a 12:12 h light/dark cycle, and food and water were available *ad libitum*. Mice were assigned to experimental groups using a random number generator.

For rat experiments, Sprague-Dawley rats were purchased from Taconic to arrive weighing approximately 250g each. Once rats arrived in the facility, they were habituated for a minimum of 5 days. Prior to experiments animals were handled for a minimum of 15 minutes/day over 3 days. Animals were housed in same-sex groups of 2 on a 12:12 h light/dark cycle with access to food and water *ad libitum*. All rats were assigned to their experimental group using a random number generator, with no more than 1 animal per experimental group in each housing cage. Experimenters were blinded to treatment groups for all behavioral experiments.

### Drugs

Recombinant soluble human interleukin 6 receptor (IL-6r) and recombinant human IL-6 (IL-6) were obtained from R&D systems (Minneapolis, MN). We have previously shown that these IL-6 signaling components produce equivalent effects in hyperalgesic priming (Paige et al., 2018). Prostaglandin E_2_ (PGE_2_) was obtained from Cayman Chemical Company (Ann Arbor, MI). Rat ∝-CGRP_8-37_ and rat α-CGRP were obtained from Bachem (Torrance, CA) or Tocris (Minneapolis, MN). CLP257 was ordered from Tocris. IL-6 and IL-6r stock solutions were made in sterile 1X PBS and did not undergo multiple freeze/thaw cycles. PGE_2_ stock solution was made in 100% ethanol. Rat ∝-CGRP_8-37_ and rat α-CGRP were made in sterile filtered 1X PBS for *in vivo* experiments and in ddH_2_O for electrophysiological recordings. Monoclonal mAbs were shipped to UT Dallas from Alder Pharmaceuticals (Seattle, WA – Alder has now been acquired by Lundbeck) at the working concentrations for injection. The vehicle for both the CGRP mAb and control mAb was 25 mM Histidine and 250 mM Sorbitol at pH 6.0. All drugs were diluted to the final concentration in sterile filtered 0.9% saline and kept on ice until immediately prior to the injections.

### von Frey Testing

Prior to testing, animals were placed in an acrylic box with mesh flooring and allowed to acclimate for a minimum of 1 h on the day of testing. Baseline paw withdrawal thresholds were determined before any experimental work. During experiments, if time points were earlier than 1h post treatment injection, animals were allowed to habituate for 1h, injections were given, and animals were placed back into the acrylic boxes. Paw withdrawal threshold was then determined using calibrated von Frey filaments using the up-down method (Chaplan et al., 1994).

### Hyperalgesic Priming

In order to establish hyperalgesic priming, 0.1 ng of IL-6 or IL-6r was injected intraplantarly (I.Pl.) into the left hindpaw, as noted for each experiment. Paw withdrawal threshold was then measured at 3, 24, 72 h, and then 7 d or until mice returned to their baseline mechanical withdrawal threshold. For experiments in rats, animals were tested every other day after the 72 h timepoint until they had returned to baseline. Animals were then injected with 100 ng of PGE_2_ and mechanical paw withdrawal threshold was measured at 3 and 24 h after injection.

### Spared Nerve Injury (SNI)

Animals were anesthetized using 4% isoflurane gas and kept on a heating pad for the entire length of surgery. A small incision was made in the left leg and the tibial and common peroneal branches of the sciatic nerve were ligated and cut (Decosterd and Woolf, 2000). The sural nerve was left intact. The incision was closed using 2 staples and animals were given a subcutaneous injection of 1 mg/mL Gentamicin. Mice were returned to their home cage and were tested at 21 days post-surgery to determine if mechanical hypersensitivity had developed. Drug testing was done after animals were confirmed to show mechanical hypersensitivity in the affected paw.

### Brennan Incision Model

Mice were anesthetized using 4% isoflurane gas. An incision was made using a scalpel through the skin and underlying fascia of the left hindpaw. Two sutures were used to close the incision and animals were allowed to recover in their home cages for 24h before any behavioral testing was performed (Banik et al., 2006).

### Intrathecal Injections

Animals were anesthetized using isoflurane gas – 4% for induction, 1.5% for maintenance anesthesia. Injections were performed as described by Hylden and Wilcox (Hylden and Wilcox, 1980). A total volume of 5 *μ*L was injected for each drug using a 50 *μ*L Hamilton syringe with attached ½ in. 30-gauge needle. Animals were allowed to recover in their home cage for a minimum of 10 minutes before any behavioral testing was performed.

### Immunohistochemistry Analysis

#### Tissue preparation and immunohistochemistry

Mice were anesthetized with xylazine (12 mg/kg) and ketamine (80 mg/kg) administered I.P. Animals were then transcardially perfused with 10% PFA in 0.1 M PB at a pH of 7.4. Spinal cords were dissected and post-fixed in 4% PFA for 2h at RT. Fixed spinal cords were then moved to a 25% sucrose suspension at 4° C for a minimum of 24 h. These spinal cords were then stored in an anti-freeze solution at −20° C until processed for immunohistochemistry.

Mice spinal cord explants were obtained by laminectomy from ketamine/xylazine anesthetized C57BL6/N male and female mice. The whole spinal cord was immersed in ice-cold S-ACSF and the lumbar enlargement was isolated and divided in two pieces (rostral and caudal). Each section was left to recover in ACSF at 34°C for 30 min. Consecutively, the two lumbar pieces from each animal were transferred separately to room temperature ACSF supplemented with 50 nM CGRP or vehicle. Rostral-caudal fractions were randomly assigned to the treatments to prevent differences arising from spinal segment variability. After 3h, spinal cords were fixed in 4% PFA for 20h at 4°C. Fixed spinal cords were moved to 25% sucrose solution at 4°C for a minimum of 24h.

The fixed spinal cord tissue was sectioned transversally using a Leica VT1200S vibratome in 50 μm thickness slices. Sections were permeabilized in PBS (pH 7.4) with 0.2% Triton (PBST) for 10 min, washed twice in PBS and incubated overnight at 4°C in primary anti-KCC2 raised in rabbit (1:1000, Millipore/Upstate, Cat. #07–432), anti-CGRP raised in mouse (1:5000, Sigma Cat. #C7113) and anti-NeuN raised in chicken (1:1000, EMD Millipore-Sigma Cat. #6B9155) antibodies diluted in PBST containing 10% normal goat serum. After washing in PBS, the sections were incubated for 2 h at room temperature in a solution containing a mixture of goat-Cy3 anti-rabbit (1:500, Jackson ImmunoResearch Laboratories, Cat. #111–165–144), goat anti-chicken Alexa 647 (1:500, Thermo Fisher Scientific Cat. #A-21449), goat anti-mouse Alexa 405 (1:500, Thermo Fisher Scientific Cat. #A-31553) and IB4 conjugated with Alexa 488 (1:500, Thermo Fisher Scientific Cat. #I21411) diluted in PBST (pH 7.4) containing 10% normal goat serum. Upon completion of this step, sections were washed three times with PBS and then were mounted on Superfrost Plus glass slides (Thermo Fisher Scientific catalog #10149870) and cover-slipped (Catalog #12-544-E) using fluorescence mounting medium (Dako, Cat. #S3023).

#### Confocal microscopy imaging

Images were captured using a confocal microscope (Zeiss, LSM 700; Oberkochen, Germany). Appropriate filters were selected for the separate detection of Alexa 405, Alexa 488, Cy3 and Alexa 647, using a multi-track scanning method. Eight-bit images were taken with a x63/1.4 oil-immersion objective lens. To ensure consistency among samples, all parameters of laser power, pinhole size and image detection were kept unchanged between the image acquisitions of different samples. The chosen parameters were set so that the detection of the staining was maximal while avoiding pixel saturation. For the subsequent quantification, the channels corresponding to each staining were exported separately as TIFF files (grey scale). The channel corresponding to KCC2 staining was further analyzed using a custom-made MATLAB code.

#### Image analysis and KCC2 quantification

To calculate the intracellular and membrane KCC2 levels a custom-made MATLAB code was developed for the analysis, as previously described (Dedek et al., 2019; Ferrini et al., 2020; Lorenzo et al., 2020). Briefly, the homemade MATLAB routines used for analyzing the membrane and intracellular KCC2 provides a profiling plot of the membranes of selected neurons within the region of interest (lamina I and lamina II based on the CGRP and IB4 staining respectively). The peak of the plot defines the membrane (position 0). The values on the left-hand side of the position 0 on the X axis stand for the extracellular space while the values on the right-hand side of the position 0, stand for the intracellular space. For the final plotting, to obtain one curve per experimental group, the KCC2 values per position per animal belonging in the same group were merged. To obtain the data for the membrane KCC2 we grouped the values at position 0 for each group.

### Electrophysiology

#### Tissue preparation

C57BL6/N mice were deeply anesthetized with ketamine/xylazine. After decapitation, the vertebral column was swiftly removed and immersed in ice-cold oxygenated (95% O_2_, 5% CO_2_) sucrose-based artificial cerebrospinal fluid (S-ACSF) containing (in mM): 252 sucrose, 2.5 KCl, 2 MgCl_2_, 2 CaCl_2_, 1.25 NaH_2_PO_4_, 26 NaHCO_3_, 10 glucose, and 5 kynurenic acid. A laminectomy was performed in ice-cold S-ACSF to extract the spinal cord. The lumbar enlargement was isolated and parasagittal slices (300 µm-thick) were obtained in the same solution with a Leica vibratome. Slices were allowed to recover for 30 min at 34°C in oxygenated ACSF containing (in mM): 126 NaCl, 2.5 KCl, 2 MgCl_2_, 2 CaCl_2_, 1.25 NaH_2_PO_4_, 26 NaHCO_3_, 10 glucose. Slices were then moved to room temperature oxygenated ACSF supplemented with 50 nM of rat CGRP (Tocris) or vehicle and maintained in this solution for a minimum of 2h.

#### Cl^−^ extrusion measurements

After incubation, slices were transferred to a recording chamber where they were continuously perfused (2-3 mL/min) with oxygenated ACSF containing 1 µM tetrodotoxin (TTX; Alomone Lab), 1 µM strychnine (Sigma), 10 µM 6-cyano-7-nitroquinoxaline-2,3-dione (CNQX; Sigma) and 40 µM D(−)-2-amino-5-phosphonovaleric acid (APV; Tocris) and 50 nM of rat CGRP or vehicle. Neurons were identified using 40x water immersion-objectives. Neurons from laminae I and II were selected for the experiment, maintaining an approximate ratio of 1:2, respectively. Lamina I was identified as a narrow dark band of grey matter with a typical reticulated appearance in the edge of the dorsal white matter, and lamina II as a wider translucent band below lamina I.

Differences in KCC2 extrusion were indirectly measured by estimation of the reversal potential of GABA_A_ receptors (E_GABA_) under a challenging intracellular Cl^−^ load as previously described (Lorenzo et al., 2020). Whole cell patch-clamp recordings were performed with a MultiClamp 700B amplifier (Molecular devices). Borosilicate pipettes (3-5 MΩ) were filled with an intracellular solution containing (in mM): 115 K-methylsulfate, 25 KCl, 2 MgCl_2_, 10 HEPES, 4 ATP-Na, 0.4 GTP-Na and adjusted with KOH to a pH of 7.2. Only cells with a stable access resistance lower than 20 MΩ and a resting potential more negative than −50 mV were included for the analysis. The GABA_A_ receptor agonist muscimol was freshly dissolved in HEPES-buffered ACSF to a concentration of 500 εM, and briefly applied (30ms, 10 psi) at increasing holding potentials (12.5 mV steps). Intervals of 30s at a holding voltage of −60 mV were allowed between puffs. Data were analysed using Clampfit 10.2 (Molecular devices) and membrane potentials were corrected off-line for liquid junction potential (8mV). GABA I-V curves were obtained from the average of 3 protocol recordings and E_GABA_ was estimated as the x-axis intercept of the derived linear equation using Prism. Two different experimenters collected the data yielding to the same result. Experimenters were blinded to the animal sex.

### Statistics

All statistical analysis was performed using Prism version 8.0 and all data are shown as mean +/− standard error of the mean (SEM). Differences between each group of animals in behavioral experiments was determined using a two-way ANOVA with Bonferroni’s post-hoc test and α = 0.05. Differences between slopes in dose response curve comparisons were determined using an ANCOVA test and α = 0.05. Differences in E_GABA_ were determined using unpaired t-test after passing normality with D’Agostino & Pearson test. Multiple Wilcoxon paired-test was used for KCC2 quantification in spinal cord explants.

## Results

### Olcegepant attenuates mechanical hypersensitivity associated with hyperalgesic priming and SNI only in female mice

We first tested the effect of intrathecal (I.T.) CGRP receptor antagonists on interleukin 6 receptor (IL-6r) induced hyperalgesic priming (Dina et al., 2008; Paige et al., 2018). In this testing paradigm, animals are first challenged with 0.1 ng IL-6r injected intraplantarly (I.Pl.) and mechanical sensitivity is measured. Following the resolution of initial mechanical hypersensitivity, 100 ng PGE_2_ is injected I.Pl. as a second stimulus. Animals that received a previous injection of IL-6r show enhanced responses to the subsequent PGE_2_ injection, demonstrating the presence of hyperalgesic priming. We administered I.T. 10 *μ*g Olcegepant, a CGRP receptor antagonist, or vehicle, immediately before IL-6r injection to assess the effect of CGRP receptor blockade on the response to IL-6r and the development of hyperalgesic priming. Olcegepant attenuated the initial mechanical hypersensitivity in response to IL-6r injection in female mice but had no effect in male mice (Fig 1A; female Olcegepant effect F(1, 42) = 31.27, p < 0.0001; time effect F(6, 42) = 34.53, p < 0.0001). I.T. Olcegepant given immediately prior to IL-6r injection did not have any effect on the development of hyperalgesic priming in either male or female mice (Fig 1A). Therefore, blocking spinal CGRP receptors at the time of the initial stimulus has an effect on acute hypersensitivity in female mice, but priming is not affected in either sex.

**Figure 1:**
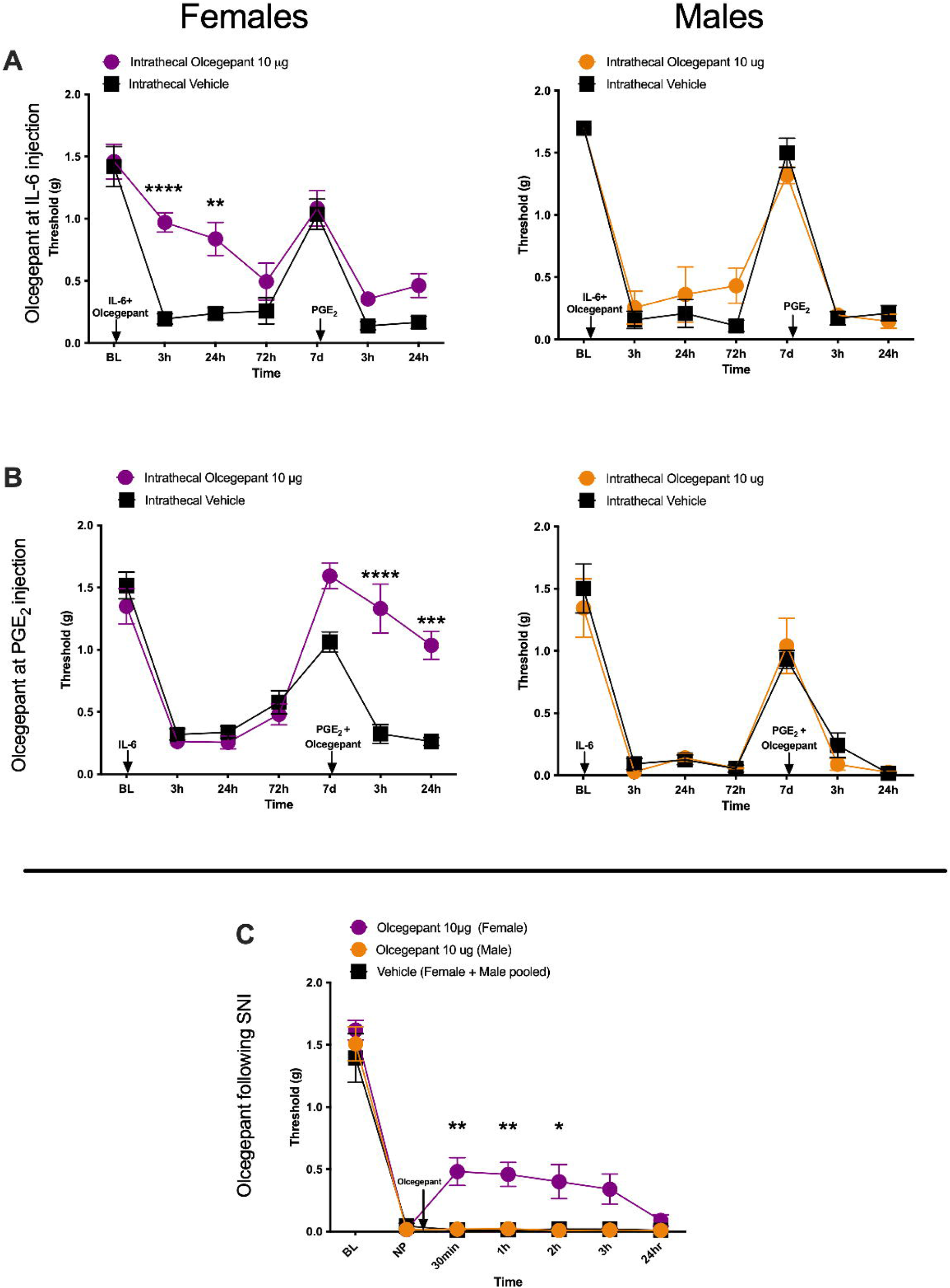
I.T. Olcegepant reduces mechanical hypersensitivity specifically in female mice. All graphs display mechanical withdrawal threshold. A. Mice received an I.T. injection of 10 *μ*g of Olcegepant prior to 0.1 ng I.Pl. IL-6r. A second, I.Pl. injection of 100ng PGE_2_ was given after initial mechanical sensitivity to IL-6r had resolved (n = 4 mice per group). B. Animals received an I.Pl. injection of 0.1 ng IL-6r. After initial mechanical hypersensitivity had resolved animals received an I.T. injection of 10 *μ*g Olcegepant and an I.Pl. injection of 100ng PGE_2_ (n = 4 mice per group). C. Animals underwent an SNI surgery. 21 d post injury, animals received a single I.T. injection of 10 *μ*g Olcegepant (n = 3 (males), 4 (females)) or vehicle (n = 5 (pooled males and females)). Differences between groups were measured using a two-way ANOVA with Bonferroni’s post hoc test, * p<0.05, ** p < 0.01, ***p < 0.001, ****p < 0.0001.

In a separate set of animals, we tested the ability of 10 *μ*g Olcegepant to reverse established hyperalgesic priming. 0.1 ng IL-6r was injected I.Pl. and mechanical hypersensitivity was measured (Fig 1B). After the initial hypersensitivity was resolved following IL-6r injection, animals received either 10 *μ*g Olcegepant or vehicle I.T. immediately prior to receiving an I.Pl. 100 ng PGE_2_ injection. Olcegepant reversed hyperalgesic priming in female mice but had no effect in male mice (Fig 1B; female Olcegepant effect F(1, 42) = 27.77, p < 0.0001; time effect F (6, 42) = 45.44, p < 0.0001).

We then tested the impact of I.T. Olcegepant in the SNI model of neuropathic pain in mice. Following the establishment of neuropathic pain, animals were given a single I.T. injection of 10 *μ*g Olcegepant or vehicle. In female mice, Olcegepant reduced mechanical hypersensitivity at 30 min, 1 h, and 2 h following injection, but the drug had no effect in male mice (Fig 1C; Olcegepant effect F (2, 63) = 25.26, p < 0.0001; time effect F (6, 63) = 123.6, p < 0.0001). These experiments demonstrate a female-specific effect of Olcegepant on mechanical hypersensitivity in hyperalgesic priming and neuropathic pain models.

### CGRP_8-37_ attenuates hyperalgesic priming, post-surgical pain, and neuropathic pain only in female mice

CGRP_8-37_ is a peptide antagonist of the CGRP receptor (Chiba et al., 1989). We chose to test a second CGRP antagonist to reduce the probability that an off-target effect explains our observations of a sex-specific effect of blocking the CGRP receptor. Animals received an I.T. injection of either 1 *μ*g CGRP_8-37_ or vehicle immediately prior to receiving an I.Pl. injection of 0.1 ng IL-6r. In female mice, CGRP_8-37_ reduced mechanical hypersensitivity induced by IL-6r and blocked the development of hyperalgesic priming as revealed by the lack of response to 100 ng PGE_2_ injection (Fig 2A; CGRP_8-37_ effect F (1, 126) = 88.54, p < 0.0001; time effect F (8, 126) = 12.30, p <0.0001). There was no effect of CGRP_8-37_ in male mice (Fig 2A). In our next experiment, 0.1 ng IL-6r was injected I.Pl. to establish priming. When the initial hypersensitivity had resolved, 1 *μ*g CGRP_8-37_ was injected I.T. immediately prior to I.Pl. 100 ng PGE_2_. Again, in female mice CGRP_8-37_ was able to reverse established hyperalgesic priming but had no impact on male mice (Fig 2B; CGRP_8-37_ effect in females F (1, 126) = 41.25, p < 0.0001; time effect in females F (8, 126) = 45.44, p <0.0001).

**Figure 2:**
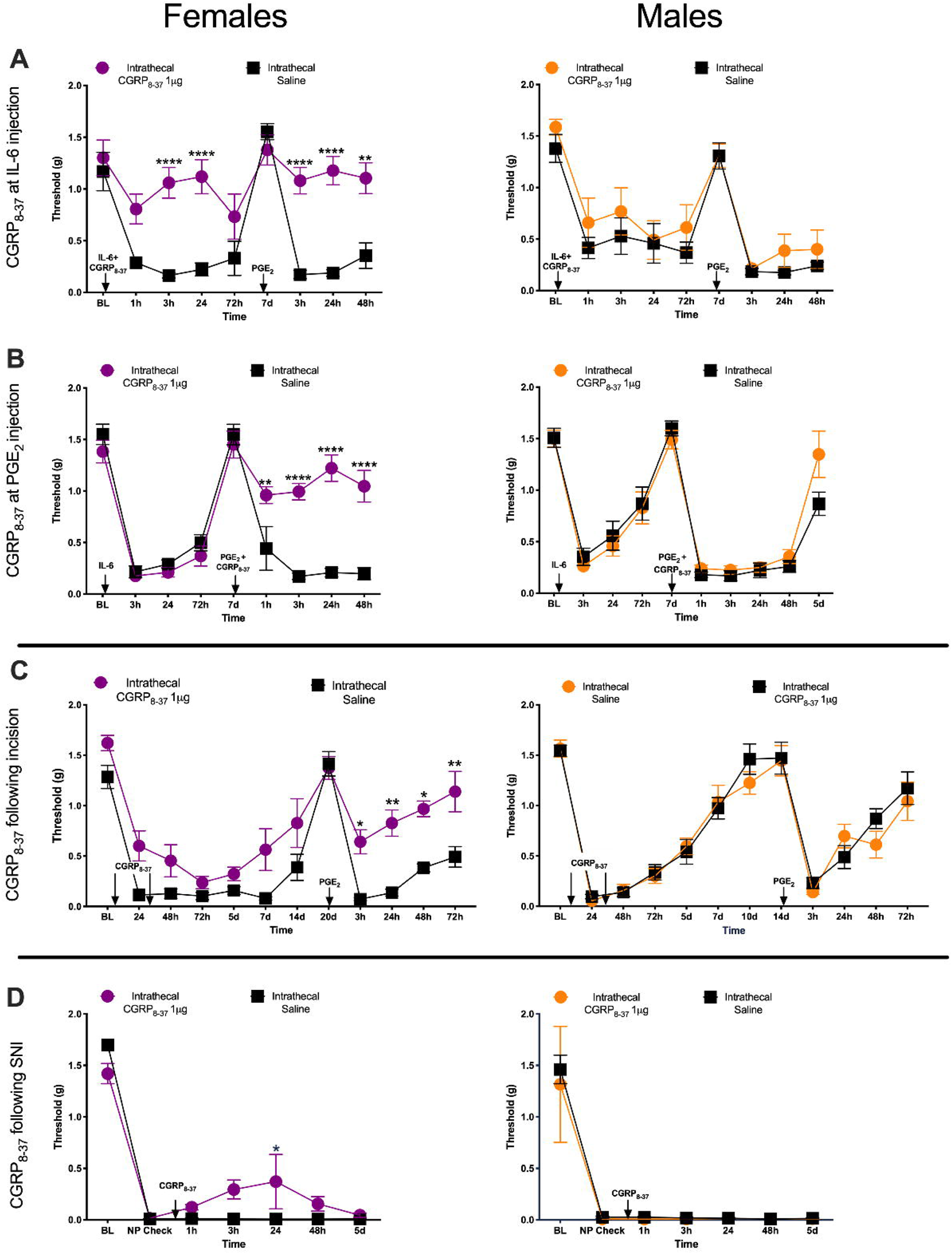
I.T. CGRP_8-37_ reduces mechanical hypersensitivity specifically in female mice in 3 pain models. A. Animals received a single I.T. injection of 1 *μ*g of CGRP_8-37_ immediately prior to an I.Pl. injection of 0.1 ng of IL-6r. Once initial hypersensitivity to I.Pl. IL-6r had resolved, animals received a second I.Pl. injection of 100 ng PGE_2_ (n = 8 mice per group). B. Animals received an I.Pl. injection of 0.1 ng IL-6r and then 7 days later an injection of 100 ng PGE2. Immediately prior to the PGE_2_ injection animals received in I.T. injection of 1 *μ*g CGRP_8-37_ (n = 7-8 mice per group). C. Animals were given a hindpaw incision and an I.T. injection of 1 *μ*g CGRP_8-37_ at the time of incision and then 24 h post incision. After the initial hypersensitivity following the incision had returned to baseline levels, animals received an I.Pl. injection of 100 ng of PGE_2_ to test for hyperalgesic priming (n = 5-6 mice per group). D. Mice were given an SNI surgery and then a single I.T. injection of 1 *μ*g CGRP_8-37_ 21 days after initial injury (n = 3 (males) and 4 (females) mice per group; NP = neuropathic pain). Differences between groups were measured using a two-way ANOVA with Bonferroni’s post hoc test, * p<0.05, ** p < 0.01, ***p < 0.001, ****p < 0.0001.

We then tested to see if the female specific effects of a CGRP antagonist would generalize to hyperalgesic priming established using a single hindpaw incision instead of an IL-6r injection. Animals were given a single incision in the hindpaw and received 2 sutures. At the time of surgery animals received a 1^st^ I.T. injection of 1 *μ*g CGRP_8-37_ or vehicle, and then a 2^nd^ injection of 1 *μ*g CGRP_8-37_ 24 h post-surgery. We gave two drug treatments in this incision-induced priming paradigm because our previous studies indicated that two treatments are necessary to block hyperalgesic priming in this model (Tillu et al., 2012). This is consistent with the duration of ongoing activity that is induced in nociceptors by the incision (Xu and Brennan, 2010). I.T. CGRP_8-37_ did not have a significant impact on mechanical hypersensitivity following the incision in either sex, but development of hyperalgesic priming was partially attenuated in female but not male mice (Fig 2C; CGRP_8-37_ effect in females F (1, 108) = 64.40, p < 0.0001; time effect in females F (11, 108) = 24.54, p < 0.0001).

We then tested the effect of I.T. CGRP_8-37_ in the SNI model. Following the establishment of neuropathic pain, on day 21 after injury, animals received a single I.T. injection of 1 *μ*g CGRP_8-37_ or vehicle. In female mice, CGRP_8-37_ had a small, but significant effect on mechanical hypersensitivity at 24 h post injection, but there was no effect at any timepoint in male mice (Fig 2C; CGRP_8-37_ effect in females F (1, 42) = 4.762, p = 0.035; time effect in females F (6, 42) = 93.18, p < 0.0001).

### CGRP-sequestering monoclonal antibody (mAb) blocks IL-6-induced mechanical hypersensitivity and development of hyperalgesic priming in female mice and rats

CGRP receptor antagonists given I.T. can act on the spinal dorsal horn or on the DRG since the I.T. space bathes both of these tissues. We sought to determine if the sex-specific CGRP antagonist effect was mediated in the peripheral or central nervous system (CNS) using CGRP sequestering mAb that do not cross the blood brain barrier (BBB) (Edvinsson, 2015). The CGRP mAb was given I.P. 24 h prior to human IL-6. These antibody experiments were performed in both mice and rats to determine if there was a species-specific effect of the CGRP mAb. We chose to use IL-6 rather than IL-6r for these experiments because while we have shown that IL-6 and IL-6r produce equivalent effects in hyperalgesic priming in mice (Paige et al., 2018), only IL-6 has been tested in rats.

In mice, animals were given an intraperitoneal (I.P.) injection of either 20 mg/kg CGRP mAb, 20 mg/kg control mAb, or mAb vehicle (see methods). Twenty-four h post I.P. injections, animals then received I.Pl. injections of either 0.1 ng IL-6 or vehicle. Following resolution of initial hypersensitivity, animals received a second injection of control or CGRP mAb 24 h prior to 100 ng I.Pl. PGE_2_. In female mice, the double CGRP mAb treatment blocked mechanical hypersensitivity following I.Pl. IL-6 and hyperalgesic priming (Fig 3A; CGRP mAb effect F (2, 15) = 68.23, p < 0.0001; time effect F (2.7, 40.4) = 43.20, p < 0.0001). However, when CGRP mAb was only given prior to PGE_2_ injection but not at the time of IL-6 injection, there was no effect on hyperalgesic priming (Fig 3A). In males there was no effect of CGRP mAb (Fig 3A).

**Figure 3:**
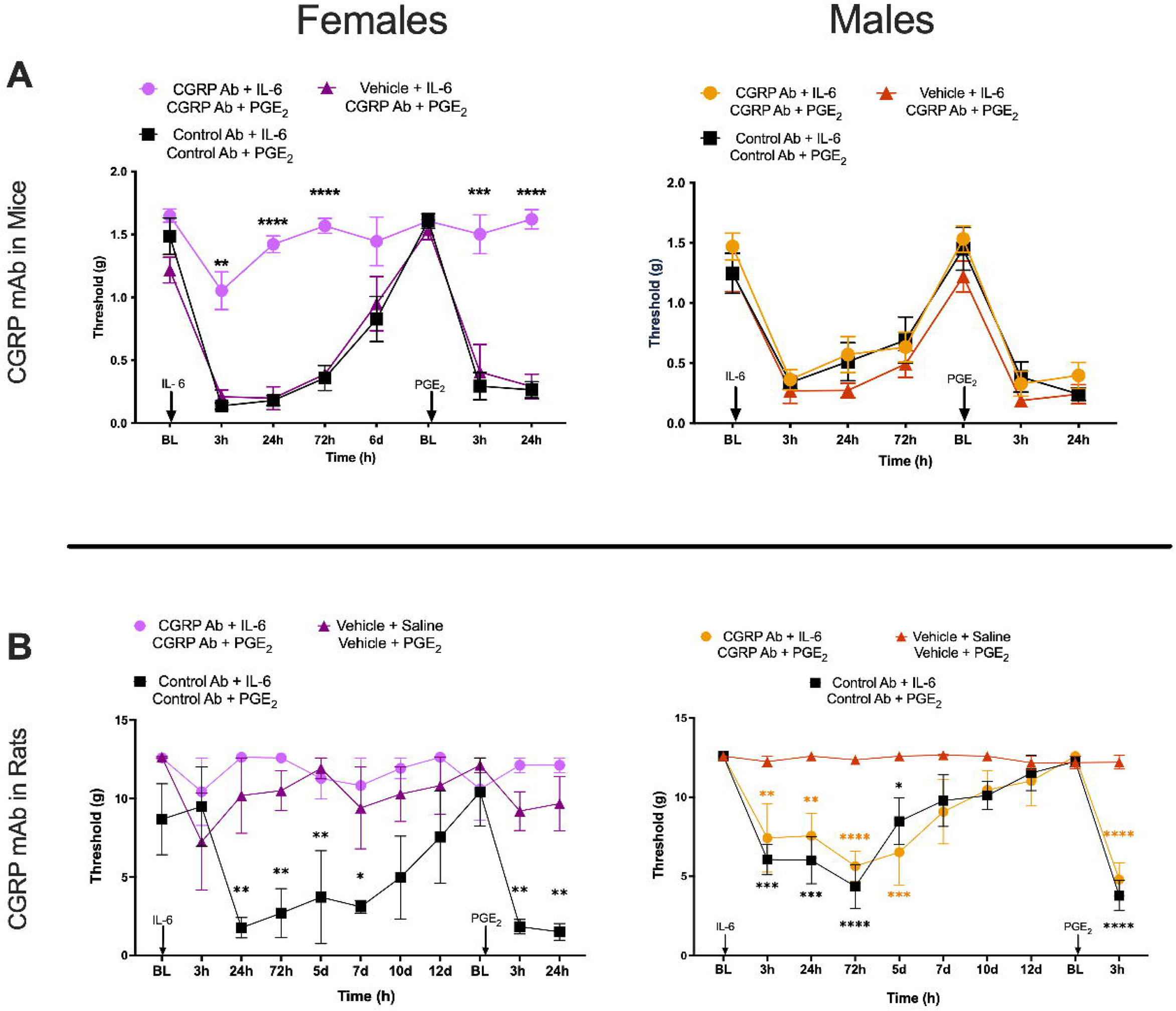
CGRP mAb block IL-6-induced mechanical hypersensitivity and development of hyperalgesic priming in female mice and rats. A. Mice were given an I.P. injection of 20 mg/kg CGRP mAb, 20 mg/kg control mAb, or vehicle 24 h prior to receiving an I.Pl. injection of 0.1 ng IL-6. After initial hypersensitivity had resolved, animals received an I.P. injection of either CGRP mAb or control mAb at the doses described above 24 h hour to a 100 ng PGE_2_ I.Pl. injection (n = 6 mice per group). Stars show significant differences vs. control Ab. B. Rats received an I.P. injection of 20 mg/kg CGRP mAb, 20 mg/kg control mAb, or vehicle 24 h prior to an I.Pl. injection of 0.1 ng IL-6 or saline and once mechanical hypersensitivity had resolved animals received an I.P. injection of the same drug or vehicle 24 h prior to I.Pl. 100ng PGE2 (n=6 rats per group). Stars show significant differences vs. vehicle treatment, color-coded by group. Differences between groups were measured using a two-way ANOVA with Bonferroni’s post hoc test, * p < 0.05, ** p < 0.01, ***p < 0.001, ****p < 0.0001.

To eliminate the possibility of a mouse-specific effect, we tested the CGRP mAb in male and female rats using the same stimulus and hyperalgesic priming paradigm. Animals received an I.P. injection of 20 mg/kg CGRP mAb, 20 mg/kg control mAb, or vehicle 24 h prior to I.Pl. injection of 0.1 ng IL-6. Following resolution of mechanical hypersensitivity, animals received a second I.P. injection of either CGRP mAb, control mAb, or vehicle 24 h prior to an I.Pl. injection of 100 ng PGE_2_. CGRP mAb blocked IL-6-induced mechanical hypersensitivity in female rats in a similar manner as we observed in female mice (Fig 3B; treatment effect F (2, 9) = 13.9, p = 0.0018; time effect F (10, 90) = 2.35, p = 0.016)). There was no effect of CGRP mAb in males (treatment effect F (2, 9) = 14.0, p = 0.0017; time effect F (9, 81) = 13.34, p = 0.0001). Therefore, in female mice and rats, CGRP mAb treatment blocks IL-6 induced mechanical hypersensitivity and hyperalgesic priming. Because receptor antagonists were effective at both stages, these findings suggest that initial effects of CGRP in these models may be peripherally-mediated while the later effects on maintenance of hyperalgesic priming are potentially CNS driven.

### Intrathecal CGRP causes prolonged mechanical hypersensitivity only in female mice

We next gave I.T. CGRP to both male and female mice. Previous work has demonstrated that I.T. CGRP in male mice and rats causes transient pain hypersensitivity but it has not been directly compared to effects in females (Cridland and Henry, 1988, 1989; Rogoz et al., 2014; Yokai et al., 2016). In the trigeminal (TG) system in rodents, CGRP-induced pain promoting effects are far more potent, and efficacious in female animals (Avona et al., 2019; Avona et al., 2021). Therefore, we tested increasing doses of α-CGRP (0.1, 0.3 and 1.0 nmol) given I.T. to male and female mice. In male mice, these doses of I.T. CGRP created transient hindpaw hypersensitivity that resolved quickly while these same 3 doses had a greater magnitude and prolonged effect in female mice (Fig 4A and B). Across a range of 5 CGRP doses (vehicle, 0.01 nmol, 0.03 nmol, 0.1 nmol, 0.3 nmol, and 1 nmol) with mechanical hypersensitivity measured only at 60 min after I.T. injection, CGRP showed increased potency (female EC_50_ = 0.0166 nmol, male EC_50_ = 0.0655 nmol) and efficacy (female E_max_ = 0.13 g threshold, male E_max_ = 0.64 g threshold) in female vs. male mice (Fig 4C). Differences between curve fits were determined using an ANCOVA test (p-value = 0.0013, F = (2, 39) = 7.931).

**Figure 4:**
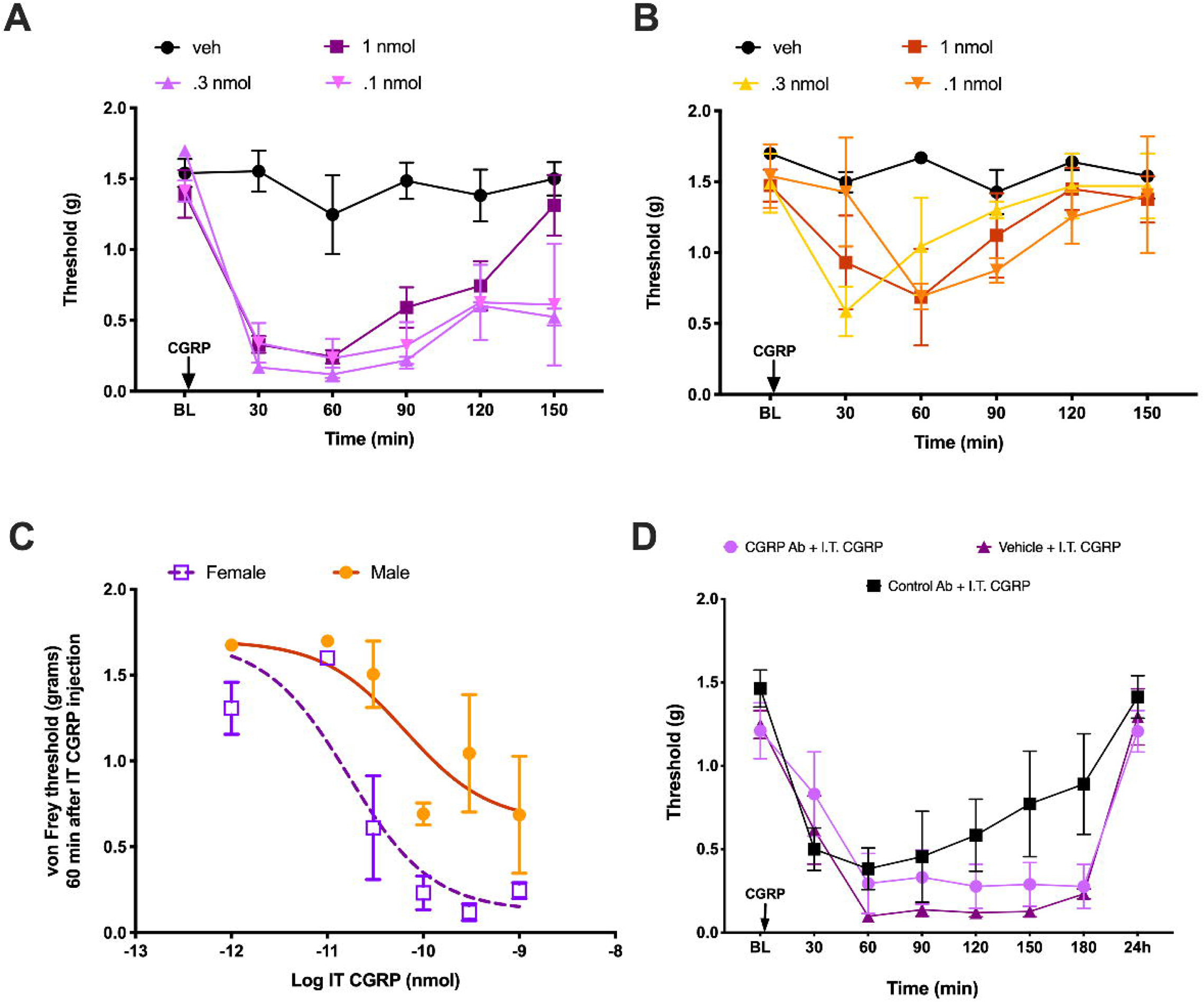
I.T. CGRP causes an increased response in female mice which is not blocked by systemic CGRP mAb. A. Female mice received an I.T. injection of CGRP in half log step increments of 0.1 nmol, 0.3 nmol, or 1 nmol of rat α-CGRP (n = 2-4 mice per dose). B. Male mice received I.T. injections of rat α-CGRP in the same doses as female mice (n = 2-4 mice per dose). C. Dose response curve of male and female mice at 60 min post I.T. CGRP. 6 increasing doses of rat α-CGRP were administered to both males and females: 0.01 nmol, 0.03 nmol, 0.1 nmol, 0.3 nmol, and 1nmol. Differences between slopes were determined using an ANCOVA test, p-value=0.0013 (F = 7.931 (Dfn = 2, Dfd = 39); n = 11 mice for female dose response curve, n = 12 mice for male dose response curve). D. Female mice received an I.P. injection of 20 mg/kg CGRP mAb, 20 mg/kg control mAb, or vehicle 24h prior to I.T. injection of 0.1 nmol rat α-CGRP and mechanical hypersensitivity was measured following these I.T. injections (n = 6 mice per group). Differences between groups were measured using a two-way ANOVA with Bonferroni’s post hoc test.

To test whether the effects of I.T. CGRP were likely centrally or peripherally mediated we gave female mice I.P. 20 mg/kg CGRP mAb, 20 mg/kg control mAb, or vehicle injection 24h prior to receiving an I.T. injection of 0.1 nmol CGRP. The systemically administered CGRP mAb was not able to block the hindpaw hypersensitivity resulting from the I.T. CGRP injection (Fig 4D). These results suggest that CGRP has increased potency and efficacy in promoting mechanical hypersensitivity in female mice via a CNS-mediated mechanism of action.

### CGRP depolarizes the GABA_A_ reversal potential (E_GABA_) in female dorsal horn neurons

The reversal potential of ionic currents gated by GABA_A_ receptors (E_GABA_) is critical for effective inhibition (Doyon et al., 2011; Doyon et al., 2016). In CNS neurons, the K^+^-Cl^−^ cotransporter KCC2 is the primary contributor to setting intracellular Cl^−^, hence E_GABA_ (Doyon et al., 2011). Previous work has demonstrated that decreased KCC2 expression and/or increased KCC2 internalization in dorsal horn neurons following peripheral nerve injury is a causative factor in mechanical hypersensitivity (Coull et al., 2003; Ferrini et al., 2013; Ferrini et al., 2020; Lorenzo et al., 2020). This mechanism is engaged in male and female rodents, however, there is a sex difference in the underlying mechanisms regulating KCC2. In males this involves BDNF signaling (Sorge et al., 2015; Moy et al., 2019), but in females the mechanism is not clearly elucidated. Since our finding suggests that CGRP affects aspects of nociceptive behavior in females by acting on central neurons (Fig. 4D), we sought to determine whether CGRP signaling regulates KCC2 extrusion capacity in the spinal dorsal horn (laminae I and II) of female and male mice. Detection of differences in KCC2 activity requires challenging the transporter, so we measured E_GABA_ in whole-cell configuration by imposing a Cl^−^ load (29 mM Cl^−^) through the recording pipette (Ferrini et al., 2020). Interestingly, while there was no difference in E_GABA_ between females and males at resting conditions, bath incubation of CGRP (50 ng/mL, > 2 h) induced a significant depolarization of E_GABA_ exclusively in females (Female E_GABA_: control = −43.68 ± 1.06 mV and CGRP = −40.54 ± 0.37 mV, unpaired t-test: t = 2.795, p = 0.011; male E_GABA_: control = −43.19 ± 0.93 mV and CGRP = −45.27 ±1.13 mV, unpaired t-test: t = 1.406, p = 0.17; Fig 5A and B).

**Figure 5:**
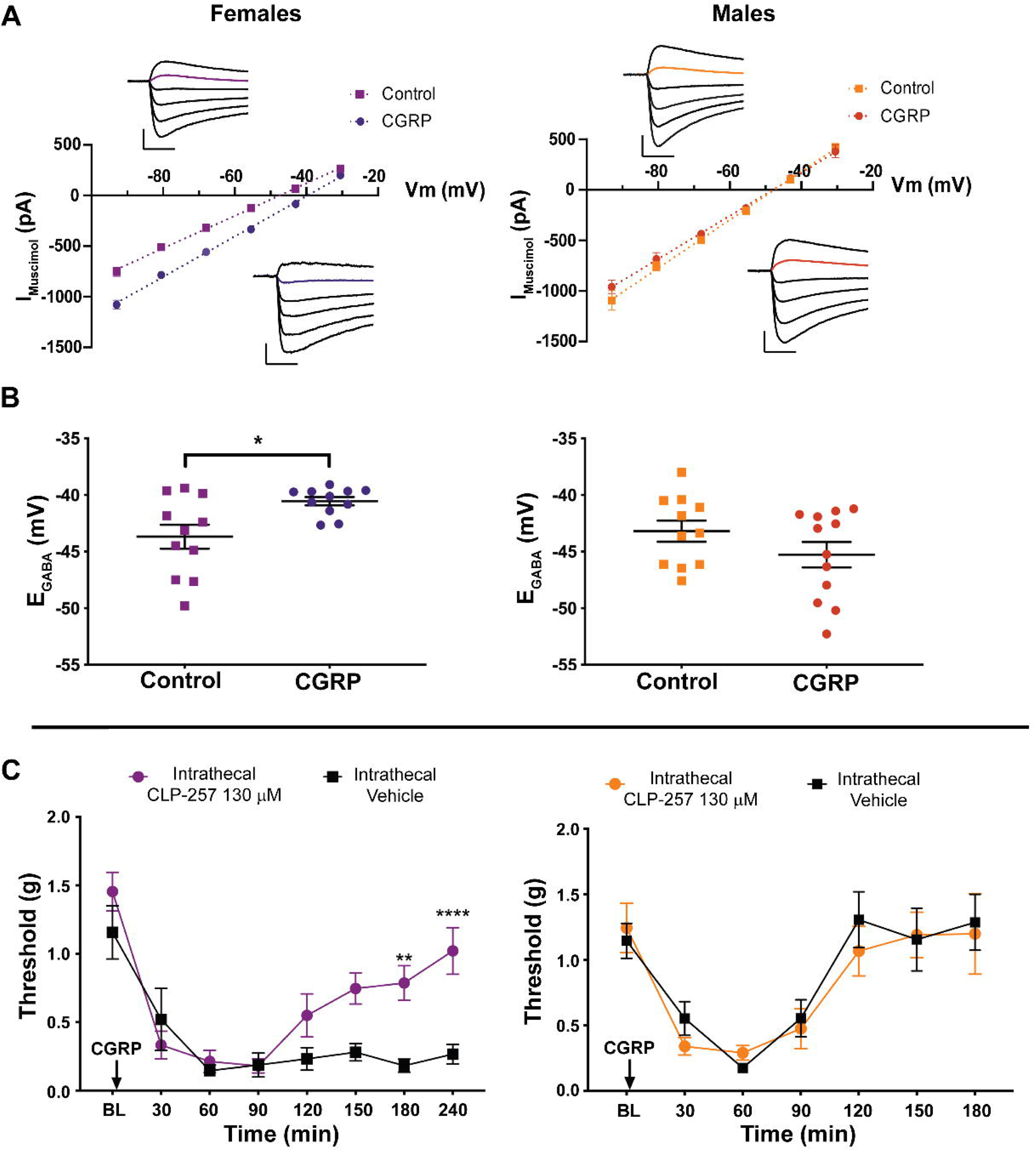
CGRP reduces Cl^−^ extrusion capacity in spinal dorsal horn neurons of female mice. A. I-V plots of a representative neurons of each experimental group. Insets: electrophysiological traces of the currents generated by 500 *μ*mol muscimol puffs at different holding voltage steps (93 mV to 30.5 mV). Scale bars: vertical = 300 pA, horizontal = 100 ms. B. KCC2 activity in laminae I and II neurons was estimated from GABA_A_ I-V curve under 29 mM Cl− load. Incubation of spinal cord slices with 50 ng/mL CGRP for more than 2h induced a depolarization of E_GABA_ only in female tissue (n=11-12 neurons per group). C. Male and female mice received an I.T. injection of 130 *μ*mol of CLP257 immediately prior to 0.1 nmol of I.T. CGRP. Mechanical hypersensitivity was measured using von Frey filament testing (n = 4 animals per group). Differences between groups were measured using a two-way ANOVA with Bonferroni’s post hoc test, **p < 0.01, *** p <0.001, ****p < 0.0001.

We then used CLP257, a KCC2 enhancer (Gagnon et al., 2013), to determine whether augmenting KCC2 activity could reverse CGRP-evoked mechanical hypersensitivity in female mice. We administered I.T. 130 *μ*mol CLP257 immediately prior to I.T. 0.1 nmol CGRP. CLP257 was able to decrease hindpaw hypersensitivity caused by I.T. CGRP in female mice, but we did not observe an effect in male mice (Fig 5C; female CLP257 effect F (1, 6) = 10.46, p = 0.018; time effect F (7, 42) = 20.36, p < 0.0001). Of note, hindpaw sensitivity caused by I.T. CGRP returned to baseline levels in male mice by the same timepoint CLP257 began to have an effect in female mice. These findings link the effect of CGRP on KCC2 to mechanical hypersensitivity in female mice.

### CLP257 inhibits hyperalgesic priming in male and female mice

To determine if KCC2 potentiation could block or reverse hyperalgesic priming, we gave male and female mice I.T. CLP257 either prior to IL-6 injection or prior to PGE_2_ in previously primed mice. I.T. CLP257 (130 *μ*mol) immediately prior to I.Pl. 0.1ng IL-6 effectively blocked IL-6 induced mechanical hypersensitivity in both sexes (Fig 6A; female CLP257 effect F (1, 6) = 42.21, p = 0.0006; time effect F (6, 36) = 17.56 p < 0.0001; male CLP257 effect F (1, 6) = 61.10, p = 0.0002; time effect F (6, 36) = 8.45 p < 0.0001). After mechanical hypersensitivity had resolved, animals received a second I.Pl. injection, this time 100ng PGE_2_. In both males and females that previously received CLP257, the development of hyperalgesic priming was blocked (Fig 6A).

**Figure 6:**
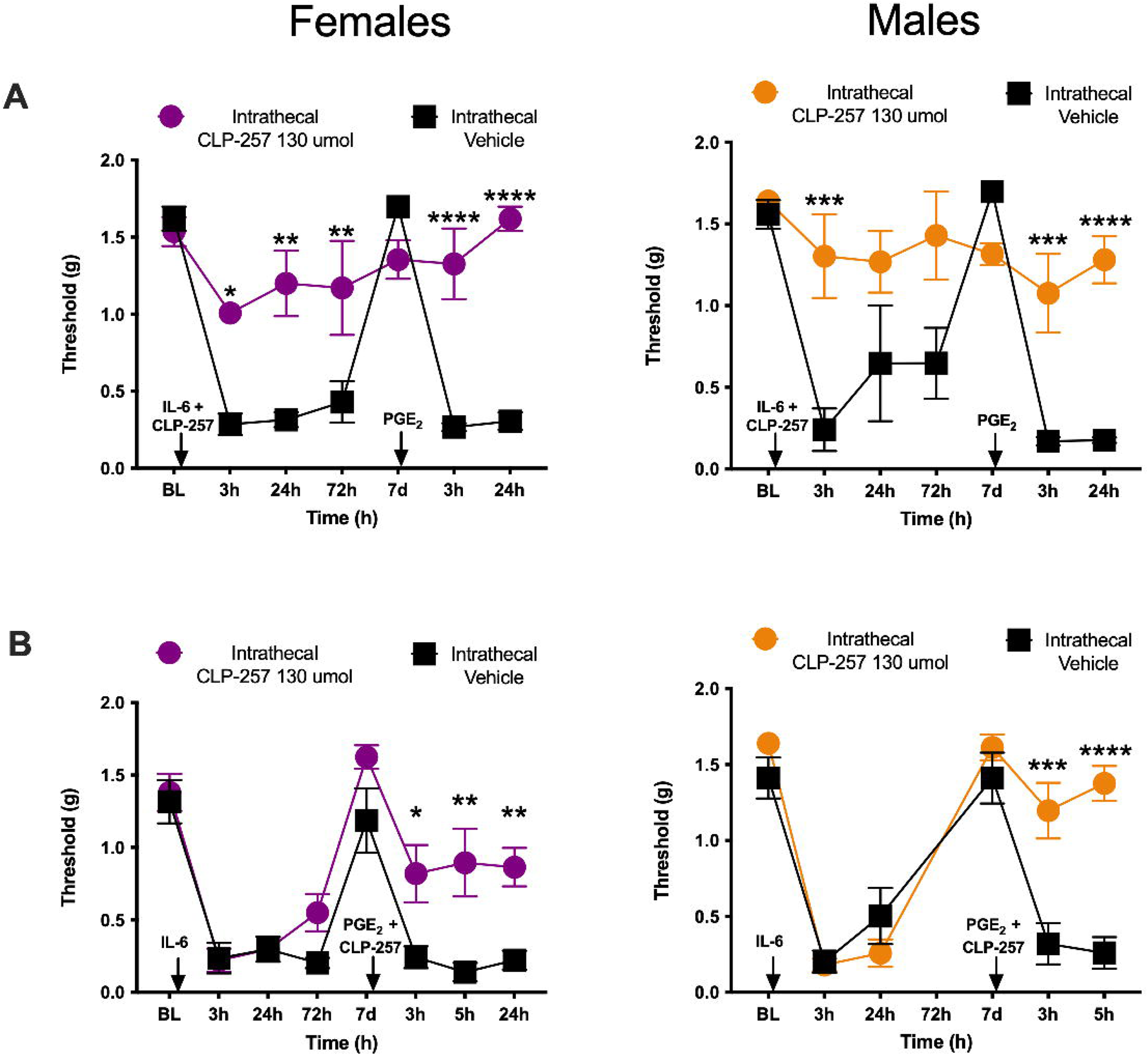
CLP257 blocks and reverses hyperalgesic priming in both male and female mice. A. Animals received an I.T. injection of 130μmol of CLP257 immediately prior to an I.Pl. injection of 0.1ng IL-6. After initial mechanical hypersensitivity to IL-6 had resolved, mice received an I.Pl. injection of 100ng PGE2. (n=4 mice per group) B. An I.Pl. injection of 0.1ng IL-6 was given to animals and immediately prior to the I.Pl. 100ng PGE2 injection animals received an I.T. injection of 130*μ*mol CLP257. (n=4 mice per group) Differences between groups were measured using a two-way ANOVA with Bonferroni’s post hoc test, * p<0.05, ** p < 0.01, ***p < 0.001, ****p < 0.0001.

In a separate set of experiments, priming was established using I.Pl. 0.1ng IL-6 and after animals had returned to baseline mechanical hypersensitivity, they received an I.T, injection of CLP257, at the same dose, immediately prior to I.Pl. 100 ng PGE_2_. CLP257 was able to reverse hyperalgesic priming in both male and female mice (Fig 6B; female CLP257 effect F (1, 6) = 12.9, p = 0.012; time effect F (7, 42) = 31.84 p < 0.0001; male CLP257 effect F (1, 6) = 80.68, p < 0.0001; time effect F (5, 30) = 36.15, p < 0.0001). These results suggest that KCC2 dysregulation is an important factor in development and maintenance of hyperalgesic priming in both mouse sexes. While our previous experiments point to CGRP as a potential causative factor in females, a distinct mechanism, such as BDNF (Moy et al., 2019), is potentially responsible for KCC2 regulation in male mice.

### CGRP does not induce KCC2 internalization in female dorsal horn neurons

E_GABA_ depolarization is usually associated with reduced membrane expression of KCC2 (Kaila et al., 2014; Ferrini et al., 2020). We therefore hypothesized that CGRP may cause KCC2 internalization in dorsal horn neurons. To test this, we gave I.T. injections of 0.1 nmol CGRP or vehicle and sacrificed animals 1 h later to remove the spinal cord. We then processed these spinal cords to examine KCC2 localization. Contrary to our hypothesis we did not note any evidence of enhanced KCC2 internalization in response to CGRP treatment in either sex (Fig 7A-F). To examine if changes in KCC2 membrane expression may occur under conditions where CGRP application can be more precisely controlled, and at a time point when the difference between male and female is most apparent, we replicated the electrophysiological conditions at which we observed a significant shift in E_GABA_. No difference was found either on KCC2 membrane expression in either female or male spinal cord explants after 3h incubation of 50 nM CGRP, indicating that CGRP did not induce KCC2 internalization in females (Fig. 8 A-C; Multiple Wilcoxon paired test: males W = 12, p = 0.46, females W = −10, p = 0.47). These results suggest that CGRP-downstream mechanisms can alter intracellular Cl^−^ and E_GABA_ through mechanisms that do not involve apparent changes in KCC2 membrane localization. This observation is consistent with recent finding that, in males, BDNF-signalling *per se*, in absence of NMDA-signalling, causes a decrease in KCC2 function without internalization (Plasencia-Fernandez et al., 2019). The ability of CLP257 to enhance KCC2 function under these conditions is also consistent with previous findings that CLP257 can enhance KCC2-mediated transport even in absence of enhanced membrane expression as shown in an overexpression assay (oocytes) where membrane KCC2 was saturated (Gagnon et al., 2013).

**Figure 7:**
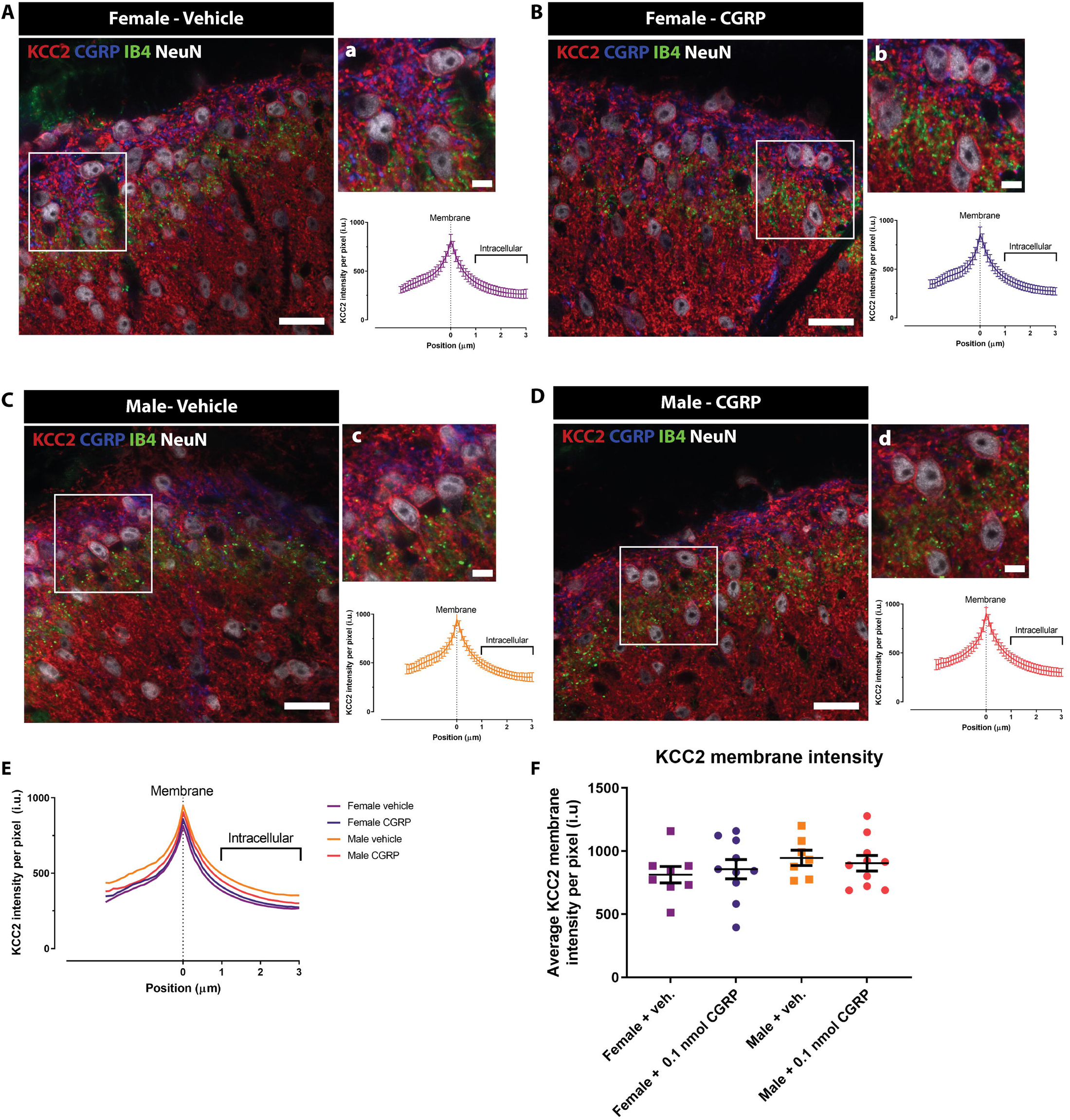
CGRP administration does not provoke KCC2 internalization in dorsal horn neurons in either female or male mice. A-D. Representative confocal images of the dorsal horn of spinal cord in female and male mice treated with either 0.1 nmol CGRP or vehicle showing CGRP, IB4, KCC2 and NeuN stainings. Scale bar in A-D = 20 *μ*m. a-d. High magnification of the areas highlighted in the white rectangle in A-D showing neurons expressing KCC2 (top) together with the average pixel KCC2 intensity plots versus distance to the membrane profile (bottom graphs). Scale bar in a-d = 5*μ*m. E. Average KCC2 intensity profiles from dorsal horn neurons of female mice treated with vehicle (violet line; n = 8 mice), female mice treated with 0.1 nmol CGRP (purple line; n = 10 mice), male mice treated with vehicle (orange line; n = 7 mice), and male mice treated with 0.1 nmol CGRP (red line; n = 10 mice). Results are presented as mean per group for visualization purposes since the curves are duplications of the graphs in a-d. F. Membrane KCC2 intensity of the four experimental groups calculated at position zero. No significant differences were observed between vehicle or CGRP treated female or male mice with Welch’s ANOVA test, n = 7 – 10 mice per group.

**Figure 8:**
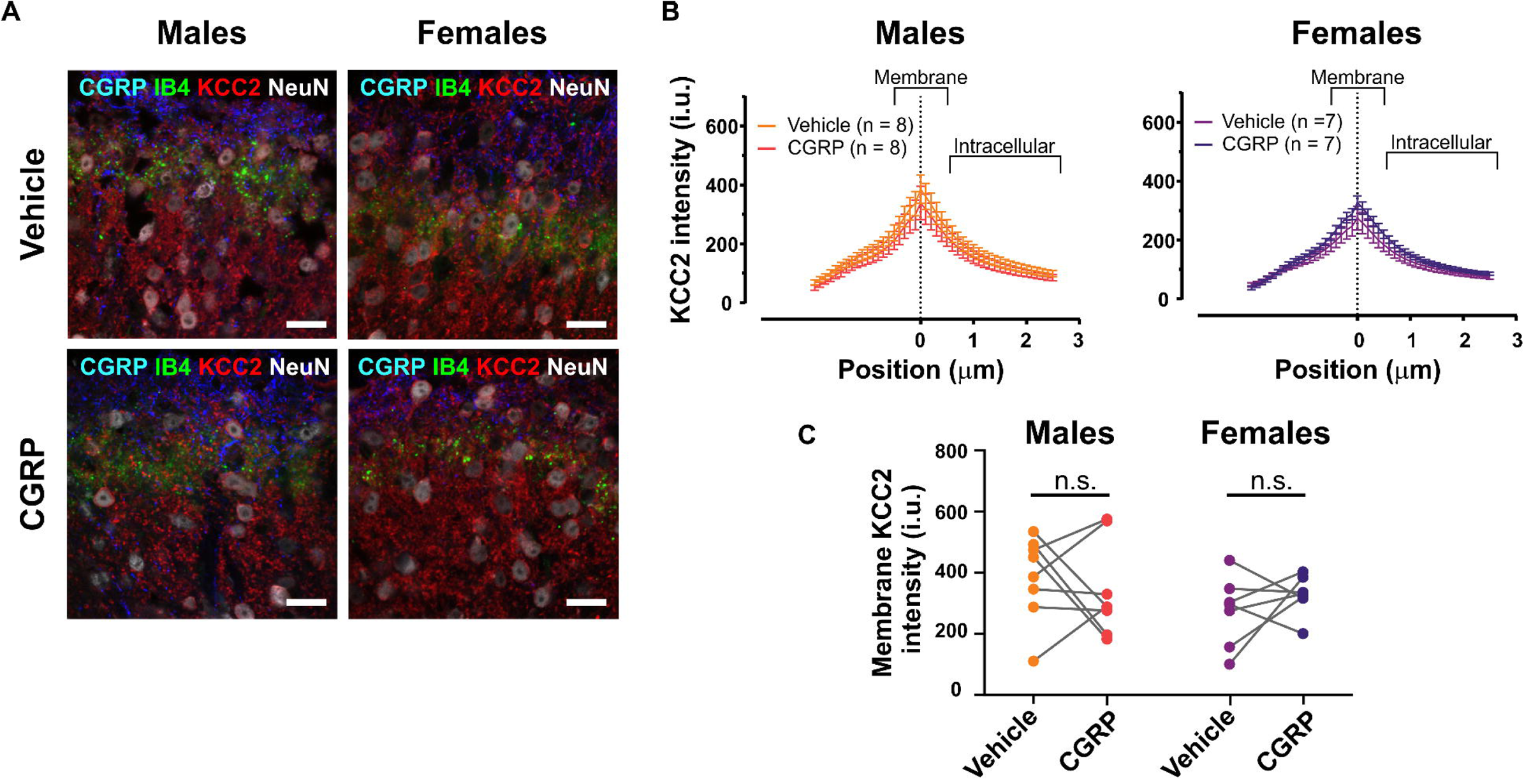
CGRP incubation for 3h does not cause KCC2 internalization in SDH neurons in either female or male mice spinal cord explants. (A) Representative confocal images of the dorsal horn of spinal cord explants from female and male mice incubated with either 50 nM CGRP or vehicle for 3h. Showing CGRP, IB4, KCC2 and NeuN stainings. Scale bar = 20 *μ*m. (B) Average KCC2 intensity profiles from dorsal horn neurons of male (left; *n* = 8 mice) and female (right; *n* = 7 mice) mice explants incubated with vehicle (orange and violet lines), or 50 nM CGRP (red and purple lines). Results are presented as mean per group. (C) Membrane KCC2 intensity of the four experimental groups calculated at position zero. No significant differences were observed between vehicle or CGRP treated female or male mice spinal cord explants.

## Discussion

Our findings support the conclusion that CGRP promotes mechanical sensitization in the DRG/spinal cord system, but primarily in female rodents. This adds to a growing body of evidence identifying signaling pathways that promote pain specifically in female animals in numerous models of chronic pain (Patil et al., 2013; Sorge et al., 2015; Avona et al., 2019; Patil et al., 2019b; Patil et al., 2019a; Paige et al., 2020; Avona et al., 2021). We found that CGRP, acting both in the periphery and CNS regulates early pain signaling in female mice and rats but that more persistent effects, such as those involved in the maintenance of hyperalgesic priming, require CNS CGRP signaling. The finding that I.T. applied CGRP-evoked mechanical hypersensitivity was more potent and efficacious in females and not blocked by a CGRP-targeting mAb peripherally injected further supports this contention. Collectively, our study raises the possibility that CGRP receptor blocking therapies that enter the CNS may be effective for non-migraine pain in women. This hypothesis can be tested in future clinical trials as the number of approved drugs targeting the CGRP system has expanded dramatically in recent years (Yuan et al., 2019).

We demonstrated that CGRP depolarizes E_GABA_ in the neurons of the dorsal horn of the spinal cord specifically in female mice. E_GABA_ is controlled by the expression and localization of KCC2 in dorsal horn neurons (Coull et al., 2003; Kaila et al., 2014; Price and Prescott, 2015; Doyon et al., 2016). Previous work has demonstrated some distinct mechanisms underlying KCC2 function in male and female mice and rats. While peripheral nerve injury causes KCC2 downregulation in both sexes (Mapplebeck et al., 2019), and KCC2 activators show efficacy in male and female rodents in neuropathic pain models, a microglial P2X4-BDNF-TrkB mediated mechanism of KCC2 downregulation is engaged only in male rodents (Mapplebeck et al., 2018; Mapplebeck et al., 2019). In our experiments, while we saw a change in E_GABA_ in female mice, we did not observe any sign of CGRP-regulated internalization of KCC2. Over the time course of our experiments, it is unlikely that transcriptional downregulation could explain the observed effect, so we did not examine KCC2 expression with quantitative PCR, but we cannot rule out such effects. It may be that a different mechanism downstream of CGRP receptors controls KCC2 function in female mice. Possibilities include phosphorylation regulated transporter function (Friedel et al., 2015; Kahle and Delpire, 2016) or targeted degradation of KCC2 pools (Plasencia-Fernandez et al., 2019). Nevertheless, the female-specific effects of CGRP appear to involve KCC2 function because its acute administration in slices effects E_GABA_ and its systemic action can be reversed by CLP257. Interestingly, and in agreement with previous studies in neuropathic pain models (Mapplebeck et al., 2019), the KCC2 enhancer CLP257 could both block and reverse the establishment of hyperalgesic priming in male and female mice. This indicates that while different upstream mechanisms governing KCC2 function are likely engaged in male and female mice, the co-transporter appears to play an important role in hyperalgesic priming in both sexes.

We addressed two issues around the pharmacology of CGRP in hyperalgesic priming in female mice and rats using distinct CGRP antagonism approaches: (1) whether CGRP antagonism could prevent and/or reverse hyperalgesic priming, and (2) whether these effects were centrally or peripherally mediated. We found that Olcegepant and CGRP mAb were only able to block hyperalgesic priming from occurring, but I.T. CGRP_8-37_ was able to both block and reverse priming in female animals. An explanation of this observation is the location in which hyperalgesic priming is mediated – with the initiation phase of priming being mediated in the DRG compartment and the maintenance phase having its locus in the CNS. I.T. injections result in drug potentially bathing both the spinal cord and the DRG whereas CGRP mAb are restricted to the periphery when injected systemically. However, only intraspinal injections, which require a laminectomy, assure that a compound will enter the spinal dorsal horn (Banks, 2006). Why did CGRP_8-37_ reverse hyperalgesic priming but olcegepant did not? One possible explanation is that olcegepant does not cross the blood brain barrier when given by an I.T. injection and therefore may only act on the DRG in the I.T. space (Tvedskov et al., 2010). On the other hand, although CGRP_8-37_ is a large, hydrophobic molecule, there is evidence that it crosses the blood brain barrier and therefore may access CGRP receptors in the dorsal horn when given I.T. (Edvinsson et al., 2007). Another possibility is that there may be separate receptors involved in these observed effects. Small molecule CGRP antagonists can bind to both CGRP and amylin receptors. CGRP receptors are composed of 2 separated subunits – the receptor activity-modifying protein 1 (RAMP1) and the calcitonin receptor-like receptor (CLR) (McLatchie et al., 1998); The Amylin receptor is composed of RAMP1 and the calcitonin receptor (CTR) (Hay et al., 2018). Both of these receptors have shown strong affinity for CGRP antagonists, which could lead to varying effects of small molecule antagonists (Hay et al., 2018). These considerations need to be weighed when designing potential clinical studies with these drugs (Yuan et al., 2019).

There are several limitations to our study. First, we acknowledge that the sample size for many of our mouse behavioral experiments are small. We designed our mouse behavioral experiments with a group size of 8 per treatment with an equal split of male and female mice. We found a striking sex difference despite the small sample size that was consistent across experiments with different CGRP receptor antagonists. Second, we have not discovered a mechanism that explains the enhanced *in vivo* potency and efficacy for CGRP in female rodents. Future studies will need to explore how this occurs. A possibility could be sex-hormone effects on receptor signaling coupling. Finally, we have not controlled for fluctuations in the sex hormone cycle in female rodents. Given that the time course of our hyperalgesic priming experiments temporally cover several cycles it is unlikely that changes in hormone concentration over the cycle affect our observations.

It is estimated that 20% of the population in the United States suffers from chronic pain, and women make up the majority of the patients seeking treatment for their pain (Dahlhamer et al., 2018). A potential explanation for this phenomenon is that currently available analgesics are ineffective in female patients because they target molecular mechanisms that are specific to males (Mogil, 2020; Shansky and Murphy, 2021). This creates a strong case for developing pain drugs that could be used specifically in each sex. An example of this would be repurposing recently approved CGRP-targeting mAbs and/or CGRP receptor antagonists, approved for the treatment of migraine (Yuan et al., 2019), a pain state that disproportionately impacts female patients (Ashina et al., 2021), for the treatment of other pain states in women. Our work suggests that small molecule CGRP receptor antagonists would be the better choice for such clinical trials because these drugs enter the CNS, while mAbs do not. Our work adds support to the hypothesis that mechanisms underlying chronic pain are at least partially distinct in males and females, and points to CGRP as a target that can be further explored for the development of female-specific analgesics.

## Acknowledgements

This work was supported by NINDS/NIH NS102161 (T.J.P and A.N.A.); NINDS/NIH NS065926 (T.J.P.); NINDS/NIH NS113457 (CP); NINDS/NIH NS104200 (A.N.A. and G.D.); funding from Alder Biopharmaceuticals (T.J.P.); CIHR FDN159906 grant and Canada Research Chair in Chronic Pain and Related Brain Disorders (YDK).

## Conflict of interest statement

All authors declare that they have no competing interests.

A.L.F. and L.F.G-M were employees of Alder Biopharmaceuticals.

## References

Ashina M, Katsarava Z, Do TP, Buse DC, Pozo-Rosich P, Ozge A, Krymchantowski AV, Lebedeva ER, Ravishankar K, Yu S, Sacco S, Ashina S, Younis S, Steiner TJ, Lipton RB (2021) Migraine: epidemiology and systems of care. Lancet 397:1485–1495.

Asiedu MN, Mejia G, Ossipov MK, Malan TP, Kaila K, Price TJ (2012) Modulation of spinal GABAergic analgesia by inhibition of chloride extrusion capacity in mice. The journal of pain: official journal of the American Pain Society 13:546–554.

Avona A, Burgos-Vega C, Burton MD, Akopian AN, Price TJ, Dussor G (2019) Dural Calcitonin Gene-Related Peptide Produces Female-Specific Responses in Rodent Migraine Models. J Neurosci 39:4323–4331.

Avona A, Mason BN, Burgos-Vega C, Hovhannisyan AH, Belugin SN, Mecklenburg J, Goffin V, Wajahat N, Price TJ, Akopian AN, Dussor G (2021) Meningeal CGRP-Prolactin Interaction Evokes Female-Specific Migraine Behavior. Ann Neurol 89:1129–1144.

Banik RK, Woo YC, Park SS, Brennan TJ (2006) Strain and sex influence on pain sensitivity after plantar incision in the mouse. Anesthesiology 105:1246–1253.

Banks WA (2006) The CNS as a target for peptides and peptide-based drugs. Expert Opin Drug Deliv 3:707–712.

Burgos-Vega CC, Quigley LD, Avona A, Price T, Dussor G (2016) Dural stimulation in rats causes brain-derived neurotrophic factor-dependent priming to subthreshold stimuli including a migraine trigger. Pain 157:2722–2730.

Chaplan SR, Bach FW, Pogrel JW, Chung JM, Yaksh TL (1994) Quantitative assessment of tactile allodynia in the rat paw. J Neurosci Methods 53:55–63.

Chiba T, Yamaguchi A, Yamatani T, Nakamura A, Morishita T, Inui T, Fukase M, Noda T, Fujita T (1989) Calcitonin gene-related peptide receptor antagonist human CGRP-(8-37). Am J Physiol 256:E331–335.

Coull JA, Boudreau D, Bachand K, Prescott SA, Nault F, Sik A, De Koninck P, De Koninck Y (2003) Trans-synaptic shift in anion gradient in spinal lamina I neurons as a mechanism of neuropathic pain. Nature 424:938–942.

Coull JA, Beggs S, Boudreau D, Boivin D, Tsuda M, Inoue K, Gravel C, Salter MW, De Koninck Y (2005) BDNF from microglia causes the shift in neuronal anion gradient underlying neuropathic pain. Nature 438:1017–1021.

Cridland RA, Henry JL (1988) Effects of intrathecal administration of neuropeptides on a spinal nociceptive reflex in the rat: VIP, galanin, CGRP, TRH, somatostatin and angiotensin II. Neuropeptides 11:23–32.

Cridland RA, Henry JL (1989) Intrathecal administration of CGRP in the rat attenuates a facilitation of the tail flick reflex induced by either substance P or noxious cutaneous stimulation. Neurosci Lett 102:241–246.

Dahlhamer J, Lucas J, Zelaya C, Nahin R, Mackey S, DeBar L, Kerns R, Von Korff M, Porter L, Helmick C (2018) Prevalence of Chronic Pain and High-Impact Chronic Pain Among Adults - United States, 2016. MMWR Morb Mortal Wkly Rep 67:1001–1006.

Decosterd I, Woolf CJ (2000) Spared nerve injury: an animal model of persistent peripheral neuropathic pain. Pain 87:149–158.

Dedek A, Xu J, Kandegedara CM, Lorenzo LE, Godin AG, De Koninck Y, Lombroso PJ, Tsai EC, Hildebrand ME (2019) Loss of STEP61 couples disinhibition to N-methyl-d-aspartate receptor potentiation in rodent and human spinal pain processing. Brain 142:1535–1546.

Dina OA, Green PG, Levine JD (2008) Role of interleukin-6 in chronic muscle hyperalgesic priming. Neuroscience 152:521–525.

Dodick DW, Goadsby PJ, Silberstein SD, Lipton RB, Olesen J, Ashina M, Wilks K, Kudrow D, Kroll R, Kohrman B, Bargar R, Hirman J, Smith J, investigators ALDs (2014) Safety and efficacy of ALD403, an antibody to calcitonin gene-related peptide, for the prevention of frequent episodic migraine: a randomised, double-blind, placebo-controlled, exploratory phase 2 trial. Lancet Neurol 13:1100–1107.

Doyon N, Vinay L, Prescott SA, De Koninck Y (2016) Chloride Regulation: A Dynamic Equilibrium Crucial for Synaptic Inhibition. Neuron 89:1157–1172.

Doyon N, Prescott SA, Castonguay A, Godin AG, Kroger H, De Koninck Y (2011) Efficacy of synaptic inhibition depends on multiple, dynamically interacting mechanisms implicated in chloride homeostasis. PLoS Comput Biol 7:e1002149.

Echeverry S, Shi XQ, Yang M, Huang H, Wu Y, Lorenzo LE, Perez-Sanchez J, Bonin RP, De Koninck Y, Zhang J (2017) Spinal microglia are required for long-term maintenance of neuropathic pain. Pain 158:1792–1801.

Edvinsson L (2015) CGRP receptor antagonists and antibodies against CGRP and its receptor in migraine treatment. British journal of clinical pharmacology 80:193–199.

Edvinsson L, Nilsson E, Jansen-Olesen I (2007) Inhibitory effect of BIBN4096BS, CGRP(8-37), a CGRP antibody and an RNA-Spiegelmer on CGRP induced vasodilatation in the perfused and non-perfused rat middle cerebral artery. Br J Pharmacol 150:633–640.

Ferrini F, Perez-Sanchez J, Ferland S, Lorenzo LE, Godin AG, Plasencia-Fernandez I, Cottet M, Castonguay A, Wang F, Salio C, Doyon N, Merighi A, De Koninck Y (2020) Differential chloride homeostasis in the spinal dorsal horn locally shapes synaptic metaplasticity and modality-specific sensitization. Nat Commun 11:3935.

Ferrini F, Trang T, Mattioli TA, Laffray S, Del’Guidice T, Lorenzo LE, Castonguay A, Doyon N, Zhang W, Godin AG, Mohr D, Beggs S, Vandal K, Beaulieu JM, Cahill CM, Salter MW, De Koninck Y (2013) Morphine hyperalgesia gated through microglia-mediated disruption of neuronal Cl(-) homeostasis. Nat Neurosci 16:183–192.

Friedel P, Kahle KT, Zhang J, Hertz N, Pisella LI, Buhler E, Schaller F, Duan J, Khanna AR, Bishop PN, Shokat KM, Medina I (2015) WNK1-regulated inhibitory phosphorylation of the KCC2 cotransporter maintains the depolarizing action of GABA in immature neurons. Sci Signal 8:ra65.

Gagnon M, Bergeron MJ, Lavertu G, Castonguay A, Tripathy S, Bonin RP, Perez-Sanchez J, Boudreau D, Wang B, Dumas L, Valade I, Bachand K, Jacob-Wagner M, Tardif C, Kianicka I, Isenring P, Attardo G, Coull JA, De Koninck Y (2013) Chloride extrusion enhancers as novel therapeutics for neurological diseases. Nat Med 19:1524–1528.

Hay DL, Garelja ML, Poyner DR, Walker CS (2018) Update on the pharmacology of calcitonin/CGRP family of peptides: IUPHAR Review 25. Br J Pharmacol 175:3–17.

Hylden JL, Wilcox GL (1980) Intrathecal morphine in mice: a new technique. Eur J Pharmacol 67:313–316.

Ji Y, Rizk A, Voulalas P, Aljohani H, Akerman S, Dussor G, Keller A, Masri R (2019) Sex differences in the expression of calcitonin gene-related peptide receptor components in the spinal trigeminal nucleus. Neurobiol Pain 6:100031.

Kahle KT, Delpire E (2016) Kinase-KCC2 coupling: Cl− rheostasis, disease susceptibility, therapeutic target. J Neurophysiol 115:8–18.

Kaila K, Price TJ, Payne JA, Puskarjov M, Voipio J (2014) Cation-chloride cotransporters in neuronal development, plasticity and disease. Nat Rev Neurosci 15:637–654.

Keller AF, Beggs S, Salter MW, De Koninck Y (2007) Transformation of the output of spinal lamina I neurons after nerve injury and microglia stimulation underlying neuropathic pain. Mol Pain 3:27.

Krukowski K, Eijkelkamp N, Laumet G, Hack CE, Li Y, Dougherty PM, Heijnen CJ, Kavelaars A (2016) CD8+ T Cells and Endogenous IL-10 Are Required for Resolution of Chemotherapy-Induced Neuropathic Pain. J Neurosci 36:11074–11083.

Kuraishi Y, Nanayama T, Ohno H, Minami M, Satoh M (1988) Antinociception induced in rats by intrathecal administration of antiserum against calcitonin gene-related peptide. Neurosci Lett 92:325–329.

Laumet G, Edralin JD, Dantzer R, Heijnen CJ, Kavelaars A (2019) Cisplatin educates CD8+ T cells to prevent and resolve chemotherapy-induced peripheral neuropathy in mice. Pain 160:1459–1468.

Li L, Chen SR, Chen H, Wen L, Hittelman WN, Xie JD, Pan HL (2016) Chloride Homeostasis Critically Regulates Synaptic NMDA Receptor Activity in Neuropathic Pain. Cell Rep 15:1376–1383.

Locke S, Yousefpour N, Mannarino M, Xing S, Yashmin F, Bourassa V, Ribeiro-da-Silva A (2020) Peripheral and central nervous system alterations in a rat model of inflammatory arthritis. Pain 161:1483–1496.

Lorenzo LE, Godin AG, Ferrini F, Bachand K, Plasencia-Fernandez I, Labrecque S, Girard AA, Boudreau D, Kianicka I, Gagnon M, Doyon N, Ribeiro-da-Silva A, De Koninck Y (2020) Enhancing neuronal chloride extrusion rescues alpha2/alpha3 GABAA-mediated analgesia in neuropathic pain. Nat Commun 11:869.

Mapplebeck JCS, Lorenzo LE, Lee KY, Gauthier C, Muley MM, De Koninck Y, Prescott SA, Salter MW (2019) Chloride Dysregulation through Downregulation of KCC2 Mediates Neuropathic Pain in Both Sexes. Cell Rep 28:590–596 e594.

Mapplebeck JCS, Dalgarno R, Tu Y, Moriarty O, Beggs S, Kwok CHT, Halievski K, Assi S, Mogil JS, Trang T, Salter MW (2018) Microglial P2X4R-evoked pain hypersensitivity is sexually dimorphic in rats. Pain 159:1752–1763.

McLatchie LM, Fraser NJ, Main MJ, Wise A, Brown J, Thompson N, Solari R, Lee MG, Foord SM (1998) RAMPs regulate the transport and ligand specificity of the calcitonin-receptor-like receptor. Nature 393:333–339.

Miletic G, Miletic V (2008) Loose ligation of the sciatic nerve is associated with TrkB receptor-dependent decreases in KCC2 protein levels in the ipsilateral spinal dorsal horn. Pain 137:532–539.

Mogil JS (2020) Qualitative sex differences in pain processing: emerging evidence of a biased literature. Nat Rev Neurosci 21:353–365.

Moreno-Ajona D, Perez-Rodriguez A, Goadsby PJ (2020) Gepants, calcitonin-gene-related peptide receptor antagonists: what could be their role in migraine treatment? Curr Opin Neurol 33:309–315.

Moy JK, Szabo-Pardi T, Tillu DV, Megat S, Pradhan G, Kume M, Asiedu MN, Burton MD, Dussor G, Price TJ (2019) Temporal and sex differences in the role of BDNF/TrkB signaling in hyperalgesic priming in mice and rats. Neurobiol Pain 5:100024.

Paige C, Maruthy GB, Mejia G, Dussor G, Price T (2018) Spinal Inhibition of P2XR or p38 Signaling Disrupts Hyperalgesic Priming in Male, but not Female, Mice. Neuroscience 385:133–142.

Paige C, Barba-Escobedo PA, Mecklenburg J, Patil M, Goffin V, Grattan DR, Dussor G, Akopian AN, Price TJ (2020) Neuroendocrine Mechanisms Governing Sex Differences in Hyperalgesic Priming Involve Prolactin Receptor Sensory Neuron Signaling. J Neurosci 40:7080–7090.

Patil M, Hovhannisyan AH, Wangzhou A, Mecklenburg J, Koek W, Goffin V, Grattan D, Boehm U, Dussor G, Price TJ, Akopian AN (2019a) Prolactin receptor expression in mouse dorsal root ganglia neuronal subtypes is sex-dependent. J Neuroendocrinol 31:e12759.

Patil M, Belugin S, Mecklenburg J, Wangzhou A, Paige C, Barba-Escobedo PA, Boyd JT, Goffin V, Grattan D, Boehm U, Dussor G, Price TJ, Akopian AN (2019b) Prolactin Regulates Pain Responses via a Female-Selective Nociceptor-Specific Mechanism. iScience 20:449–465.

Patil MJ, Green DP, Henry MA, Akopian AN (2013) Sex-dependent roles of prolactin and prolactin receptor in postoperative pain and hyperalgesia in mice. Neuroscience 253:132–141.

Plasencia-Fernandez I, Bergeron MJ, De Koninck Y (2019) TrkB receptors engage different signaling cascades regulating respectively KCC2 function, trafficking and degradation. IBRO Reports 6:S262.

Price TJ, Prescott SA (2015) Inhibitory regulation of the pain gate and how its failure causes pathological pain. Pain 156:789–792.

Rogoz K, Andersen HH, Kullander K, Lagerstrom MC (2014) Glutamate, substance P, and calcitonin gene-related peptide cooperate in inflammation-induced heat hyperalgesia. Mol Pharmacol 85:322–334.

Shansky RM, Murphy AZ (2021) Considering sex as a biological variable will require a global shift in science culture. Nat Neurosci 24:457–464.

Sorge RE et al. (2015) Different immune cells mediate mechanical pain hypersensitivity in male and female mice. Nat Neurosci 18:1081–1083.

Sun RQ, Tu YJ, Lawand NB, Yan JY, Lin Q, Willis WD (2004) Calcitonin gene-related peptide receptor activation produces PKA- and PKC-dependent mechanical hyperalgesia and central sensitization. J Neurophysiol 92:2859–2866.

Taves S, Berta T, Liu DL, Gan S, Chen G, Kim YH, Van de Ven T, Laufer S, Ji RR (2016) Spinal inhibition of p38 MAP kinase reduces inflammatory and neuropathic pain in male but not female mice: Sex-dependent microglial signaling in the spinal cord. Brain Behav Immun 55:70–81.

Tillu DV, Melemedjian OK, Asiedu MN, Qu N, De Felice M, Dussor G, Price TJ (2012) Resveratrol engages AMPK to attenuate ERK and mTOR signaling in sensory neurons and inhibits incision-induced acute and chronic pain. Mol Pain 8:5.

Tvedskov JF, Tfelt-Hansen P, Petersen KA, Jensen LT, Olesen J (2010) CGRP receptor antagonist olcegepant (BIBN4096BS) does not prevent glyceryl trinitrate-induced migraine. Cephalalgia 30:1346–1353.

Xu J, Brennan TJ (2010) Guarding pain and spontaneous activity of nociceptors after skin versus skin plus deep tissue incision. Anesthesiology 112:153–164.

Yokai M, Kurihara T, Miyata A (2016) Spinal astrocytic activation contributes to both induction and maintenance of pituitary adenylate cyclase-activating polypeptide type 1 receptor-induced long-lasting mechanical allodynia in mice. Mol Pain 12.

Yu X, Liu H, Hamel KA, Morvan MG, Yu S, Leff J, Guan Z, Braz JM, Basbaum AI (2020) Dorsal root ganglion macrophages contribute to both the initiation and persistence of neuropathic pain. Nat Commun 11:264.

Yuan H, Spare NM, Silberstein SD (2019) Targeting CGRP for the Prevention of Migraine and Cluster Headache: A Narrative Review. Headache 59 Suppl 2:20–32.

